# A Comprehensive Survey of Scoring Functions for Protein Docking Models

**DOI:** 10.1101/2023.12.29.573663

**Authors:** Azam Shirali, Vitalii Stebliankin, Jimeng Shi, Giri Narasimhan

## Abstract

The protein-protein docking problem is fundamental to our understanding of how proteins interact. It is also important because of its relevance to drug discovery and vaccine design. Docking methods typically involve two steps. The first step generates many possible candidate conformations, while the second step evaluates the candidates using some scoring function. Despite numerous experiments with many innovative scoring functions, a good scoring function for docking remains elusive. Deep learning models offer alternatives to using explicit empirical or mathematical functions for scoring protein-protein complexes. In this study, we perform a comprehensive **survey** of the state-of-the-art scoring functions by considering the most popular and highly performant approaches for scoring protein-protein complexes, evaluating their strengths and weaknesses to aid researchers in understanding progress made in this field. Our survey compares the state-of-the-art in classical and deep learning-based scoring methods.

## 1 Introduction and Related work

The field of structural biology has been advancing rapidly for the past few decades, and experimental approaches (such as nuclear magnetic resonance (NMR), X-ray crystallography, and cryogenic electron microscopy) have enabled biologists to determine the 3D structures of many proteins that have been published in Protein Data Bank [1]. However, almost all biological functions in living organisms depend on the interactions between proteins, and determining the structure of complexes resulting from these interactions is essential for drug discovery and therapeutic development. As with individual proteins, experimental approaches to determine the structures of complexes do exist but are costly and time-consuming, thus motivating the need for fast and accurate computational methods [2].

Computational docking methods predict the 3D structure of a complex using the 3D structures of individual proteins. These methods typically involve two steps. The first step is *sampling*, during which numerous candidate conformations are generated. Following this, a *scoring* step is performed to identify the conformations closest to the native structure. A good scoring function should correctly and efficiently retain the best near-native models. Advances in computing hardware have greatly improved the first step of docking and have been surveyed by Vakser et al. [3]. However, good scoring functions to accurately and efficiently differentiate native from non-native structures remain a challenge. This comprehensive survey will review the literature on *scoring functions* for protein-protein binding.

Scoring functions can be divided into four categories: 1) physics-based, 2) empirical-based, 3) knowledge-based, and 4) those based on machine learning (ML) or deep learning (DL) [4]. The introduction of *physics-based* scoring functions began with classical force field methods that calculated binding energy by summing the Van der Waals and electrostatic interactions between two proteins [4]. Scoring functions were improved by introducing solvent effects, polarization, and charge features [4]. All physics-based models have high computation costs [4].

*Empirical-based* methods estimate the binding score of a complex by summing up a series of weighted energy terms. With more straightforward scoring functions [5], they are computationally faster.

The *knowledge-based*, or statistical-potential scoring functions use the pairwise distances between atoms or residues in the two proteins and convert them into potentials through Boltzmann inversion [6]. Knowledge-based scoring functions offer a good balance between accuracy and speed [7].

With major advances in ML and DL, it is not surprising that many learning models exist for estimating scoring functions. ML and DL approaches can learn complex transfer functions that map a combination of interface features, energy, and accessible surface area terms to predict scoring functions [8].

Figure 1 illustrates these categories and some of the publications in each.

**Fig. 1.**
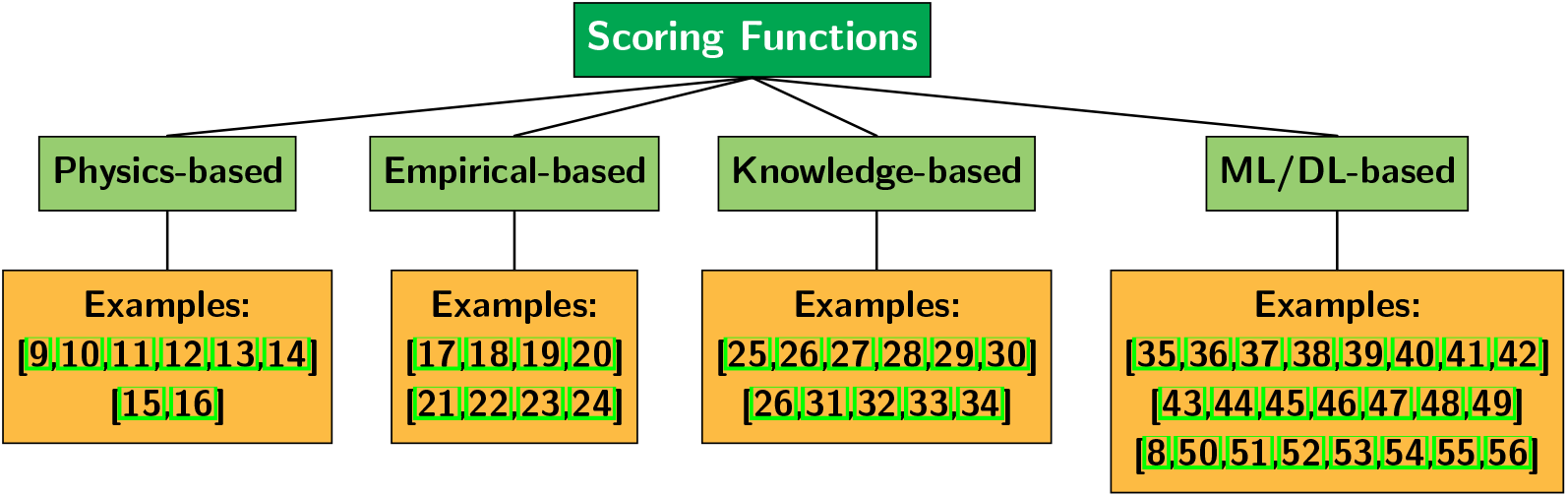
Categories of scoring functions for molecular docking models

Docking methods have been discussed in several studies and surveys. Assessment of the strengths and weaknesses of various docking methods and their range of applications are discussed in [57,58,59]. Additionally, several authors have examined the challenges and limitations of molecular docking methods, and have surveyed the shortcomings and unanswered questions within the field of molecular docking [2,60,61,62,63]. Many surveys of docking tools for drug discovery exist in the literature [64,65,66], reviewing protein-ligand docking tools. This survey will not review *docking* or protein-ligand binding, but will focus on scoring functions for protein-protein binding.

A categorization of ML- and DL-based scoring functions was presented by Li et al. [4] and reviewed by Huang et al., and Wang et al.. [7,67]. In [68], authors compared the performance of scoring functions on different types of complexes such as antibody-antigen, enzyme-inhibitor, etc. [69,70] provide a comprehensive comparison of ML-based scoring functions for virtual screening.

Moal et al. [71] evaluated 115 different scoring functions, and Su et al. [72] provided a comparative assessment of scoring functions. However, these studies did not include ML-based efforts in molecular modeling. Moreover, advancements in ML and DL for proteinprotein interaction analysis and molecular docking were reviewed recently, but without comparing them with non-ML methods [73,74]. It is essential to compare and evaluate classical and modern efforts together, not separately. Furthermore, all scoring functions were benchmarked on different datasets and have not undergone a consistent, head-to-head comparison. This raises troubling questions about whether these tools are fine-tuned and tested on specific “in-distributions” and whether they would perform as well with other “out-of-distributions” datasets [75]. Our study bridges these two gaps by conducting a comprehensive comparison across seven public and popular datasets.

The following is the organization of this paper. Section 2 summarizes all methods being compared. In section 3, we introduce the different datasets used in this work. The results of the experiments in Section 4 are followed by a discussion section.

## 2 Methods

Of the four categories of scoring functions proposed so far, the first three will be referred to as *classical* methods in the rest of the paper. This study compares the performance of seven commonly used classical methods (or their hybrids) with four cutting-edge DL-based methods. Below, we briefly summarize these methods and their properties.

### 2.1 Classical methods

**FireDock** [76] calculates the free energy change of the residues at the interface of a complex, uses an SVM to calibrate the weights, and calculates the linear weighted sum of some energy terms as the binding score. This method first refines and optimizes the binding of each candidate by allowing flexibility in the side chains and then adjusting the relative position of the protein partner using Monte Carlo minimization. Then, refined conformations are scored by calculating free energy contributions from desolvation, electrostatics, internal energies (bond stretch, angle, and torsion), hydrogen and disulfide bonds, and van der Waals interactions.

**PyDock** uses a scoring function that balances the electrostatic and desolvation energies [77]. It uses electrostatic energies with a distance-dependent dielectric constant and weighted desolvation energies for the score.

**RosettaDock**, like FireDock, reconstructs side chains by taking into account the flexibility of the chains and then performs a Monte Carlo search to refine the predicted conformations [78]. It scores the final refined complexes by minimizing an energy function that sums up contributions from van der Waals forces, hydrogen bonds, electrostatics, solvation, and side chain rotamer energies. Our experiments were conducted with PyRosetta [79], a Python-based implementation of Rosetta v3.1.

**ZRANK2** [80], an extension of ZRANK [81], calculates a linear weighted sum of energy terms representing van der Waals forces, electrostatics (attractive, repulsive, short-range, and long-range), and desolvation (using pairwise Atomic Contact Energy (ACE)). It employs RosettaDock to refine each model to generate 300 structures per predicted complex.

**AP-PISA** [82] uses a distance-dependent pairwise atomic potential in combination with a residue potential to rescore the refined complexes. The two potentials have different signals, thus increasing the chance of generating correct solutions.

**CP-PIE** [83] finds the overlap and solvent-accessible surface areas at the complex interface. They argue that the total overlap area for correct complexes is less than that for incorrect complexes, and use it as a filter to eliminate candidate complexes. The binding score also uses the number of residue contacts.

**SIPPER** [84] uses a combination of residue-residue interface propensities for each possible residue pair and a residue desolvation energy based on solvent-exposed area for scoring predicted complexes.

### 2.2 Deep Learning-based methods

Deep learning (DL) has been successfully applied across a wide range of applications and data modalities, including text [85,86], images [87,88,89], graphs [90,91], and time series data [92,93,94,95]. DL has also been applied to various problems in the field of protein science. Here we survey four DL-based approaches for scoring functions.

**GNN-DOVE** stands for Graph Neural Network (GNN)–based DOcking decoy eValuation scorE [50], and is an extension of DOVE [51]. It defines two subgraphs to represent the interface of the complex. Physicochemical features, such as the type of the atom, the number of connections, and the number of attached Hydrogen atoms, are incorporated as node features in the graph using one-hot encodings. Edges represent covalent and noncovalent bonds. A gate-augmented attention mechanism is used to learn the pattern of atomic interactions at the interface and to assess the importance of each interaction.

**DeepRank-GNN** [53] is also a GNN-based approach that learns “embeddings” of a complex. It is an extension of DeepRank [52], and represents the interface of the complex as a graph using residue-level features like type, charge, and buried surface area (BSA), for each node. Residue-level features are represented as node features. The graph representation has two subgraphs, one for the connections between residues within each protein, and the other showing the connections between residues of the two interacting proteins. The subgraphs are then passed separately to two convolution layers, one for a graph attention network (GAT) [96] and the other for edge aggregated graph attention network (EGRAT) [97]. Using these layers, the contributions of the neighbors to each node in subgraphs are calculated and weighted. The weighted sum of the neighbors’ feature representation as an aggregator helps the network to distinguish native complexes from non-native complexes.

**dMaSIF** (differentiable Molecular Surface Interaction Fingerprinting) [55], unlike its predecessor, MaSIF [54], represents the protein surface as a point cloud and determines the interactions between all atoms of the complex without using precomputed interface features. For each point of the complex interface, it calculates ten geometric features and six chemical features. These features are then passed to a sequence of (quasi-)geodesic convolutional layers for learning. The resulting binding score from dMaSIF is fast and accurate.

**PIsToN** (Protein Interfaces with Transformer Network) [56] is the most recent of the DL-based tools. PIsToN crops and isolates pairs of patches from the interface, one for each interacting protein. The information from each patch is then converted into multichannel images, with each channel representing one of the features of the interaction, including Relative Accessible Surface Area (RASA), curvature, charge, electrostatics, and hydrophobicity. PIsToN has three components. The use of a Vision Transformer (ViT) [98] differentiates it from the other methods. The embedding produced by the ViT is then combined with “hybrid” empirical energy terms, including Van der Waals, electrostatics, and desolvation, allowing it to learn the characteristics of complex binding energies. Finally, a multi-attention model groups energy terms and interface features into five groups, each containing an independent transformer network. These independent networks are aggregated into a transformer encoder to score the complexes as native or non-native decoys.

## 3 Datasets

Seven different datasets were used in this work to evaluate the methods. Links to access to these datasets are provided in Data Availability section.

**CAPRI Score** refers to the dataset used for the Critical Assessment of Predicted Interactions (CAPRI) competition that invites submissions of computational methods to compete based on a set of protein targets. Unlike the other datasets in this section, the CAPRI Score dataset was generated by a variety of different docking tools resulting in a diverse range of models, making it an excellent dataset for the evaluation of scoring functions. This dataset contains 13 targets used in rounds 13 [99] to 26 [100] with roughly 500 to 2,000 docking models per target for a total of 16,666 models [101] of varying complexity.

**CAPRI Score Refined** is a dataset produced by refining the CAPRI Score models using the HADDOCK docking tool [102]. This dataset was introduced in DeepRank [52] paper.

**Dockground** dataset 1.0 was generated using Gramm-X [103] and contains 61 complexes. Each complex has 100 non-native and at least one near-native docking model (on average, 9.83 near-native docking models per complex). There are a total of 6,725 docking models in the dataset.

**BM4** Benchmark 4.0 [104] contains 176 unbound complexes. Each protein is available in a bound and an unbound conformation. Each complex has 54,000 docking models. For each complex, we subsampled the 400 top predictions (each complex has at least one near-native docking model). The GNN-DOVE [50] study used this dataset as its training set and left 19 complexes as a test set. To make the comparison fair, we used the 19 complexes as a test set in our experiments, which provided a total of 7,600 docking models. These chosen complexes are available on Zenodo at https://doi.org/10.5281/zenodo.10442444.

**BM5** dataset contains 231 high-quality complexes. For each complex, the bonded and unbound conformations of the proteins involved in the complex are available [105]. BM5 augments BM4, which had 55 complexes. The DeepRank-GNN [53] study generated 25,300 docking models for each complex using the HADDOCK software [102]. We randomly subsampled 500 docking models for each of the 15 complexes that were not used in training or validating DeepRank-GNN, resulting in 7,500 docking models in total. These chosen complexes are available on Zenodo at https://doi.org/10.5281/zenodo.10442444.

**PDB-2023** was used in the PIsToN paper [56] and contains complexes that are deposited in the RCSB Protein Data Bank (PDB) [1] in the year 2023. All docking models were generated using HDOCK [106]. This dataset contains models for 53 heterodimers, with 126 near-native and 5174 non-native models in total.

**MaSIF dataset** was used for training and testing the MaSIF-search tool [54]. Later, the PIsToN work used 678 complexes from this dataset as a testing set, and for each complex, randomly used one non-native and one near-native conformation [56]. We used exactly this test dataset for our experiments here.

## 4 Results

In this section, we discuss the performance of all 11 methods on all 7 datasets. For DL-based methods, we used the best pretrained model, which had the best performance according to the authors’ recommendations in their corresponding paper or GitHub page. Also, we followed the settings mentioned below for each method. For GNN-DOVE we chose fold model 5 which had the best performance. For DeepRank-GNN, we used the PSSMGen [107] tool to compute Position-Specific Scoring Matrices (PSSM) features. For PIsToN, we chose a patch size of 16 Å. We calculated the average, minimum, and maximum scores for all potential contacts in dMaSIF. As the average scores had the highest AUC on all datasets, we reported them in the results. For the classical methods, we used the CCharPPI server[108], which computes the values simultaneously. No pre-calculated features and setting parameters are needed for using this server.

### Labeling the data

The CAPRI criterion was used to label all the docking models [109]. As described in [56], a docking is labeled acceptable (or correct) if:

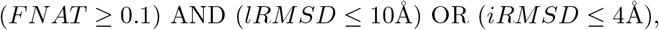

where *FNAT* is the fraction of native contacts recovered, *iRMSD* is the root mean square deviation (RMSD) between the *C_α_* atoms of the docked model and the native structure at the interface, and *lRMSD* is the global RMSD between the *C_α_* atoms of the two structures. Models for which the condition failed are labeled as incorrect [56].

### AUC ROC Measure

To evaluate the performance of the methods, the area under the curve of the receiver operating characteristic (AUC ROC) was calculated (see Figure 2). The ROC curve plots the fraction of true positives (TP) versus false positives (FP) while navigating the rankings provided by the scoring function. An ideal AUC ROC value is equal to 1, and a random 2-class classifier achieves 0.5. Additionally, various standard classification metrics are calculated, including the F1 score, precision, average precision (AP), recall, and balanced accuracy (BA) (see Supplemental Table S1).

**Fig. 2.**
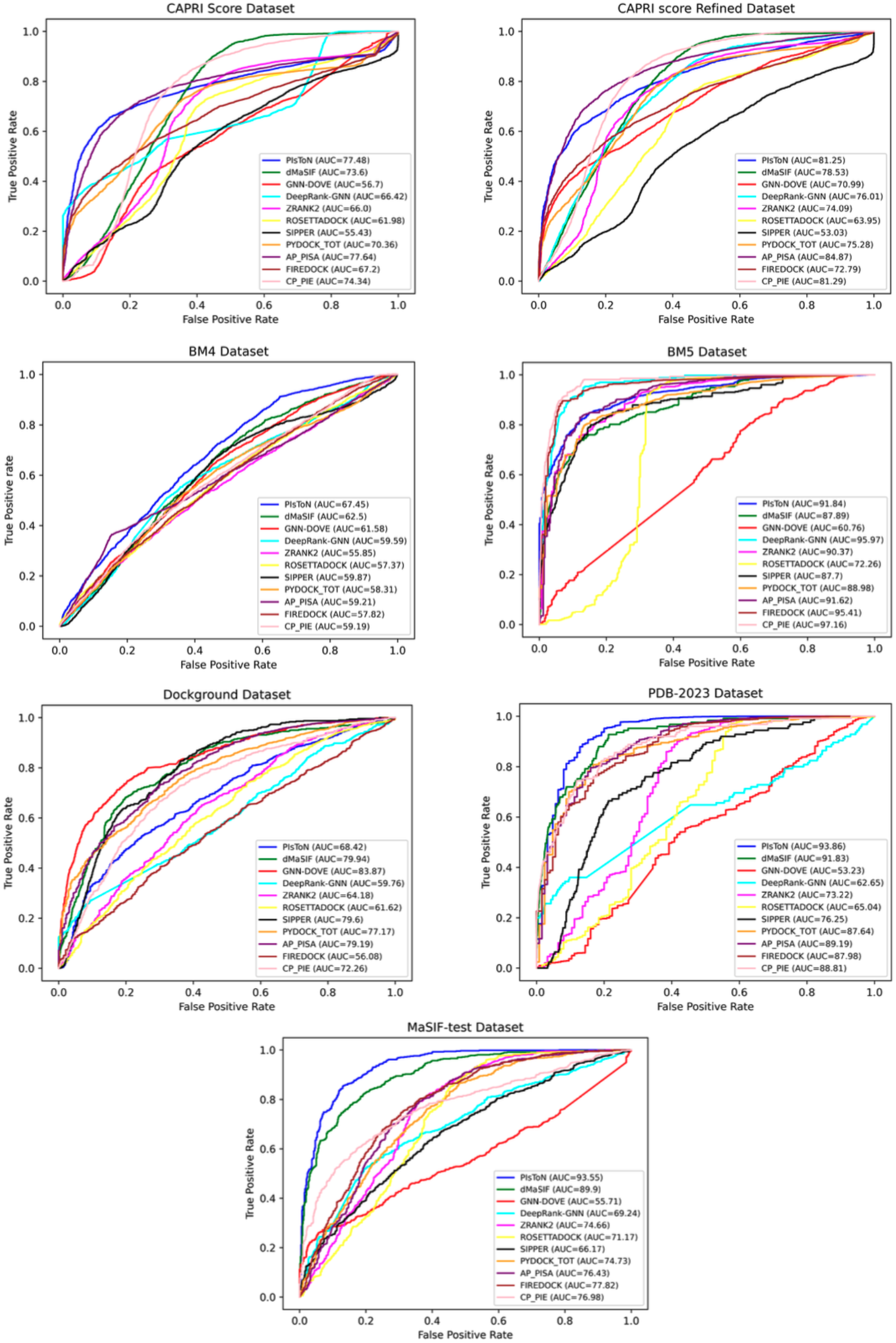
AUC ROC graphs of all eleven scoring functions on seven different datasets. The top four methods are DL-based, and the rest are classical methods (details are provided in Sec. 2)

Figure 2 shows the AUC ROC graphs of all the scoring functions on the seven datasets. In three out of seven datasets, the classical methods outperformed the DL-based methods. For the CAPRI score and the CAPRI score refined dataset, the top two performers were AP-PISA and PIsToN, with the highest AUC values. In the former, they had AUC values of 77.64 and 77.48, while in the latter, they had AUC values of 84.87 and 81.25, respectively. For the BM5 dataset, CP-PIE and DeepRank-GNN performed best with AUC values of 97.16 and 95.97, respectively. It is worth noting that DeepRank-GNN used this dataset for training and validation, but only those not in training and validation sets were used in this comparision (see Sec. 3). In short, the classical tools with the best performance in BM5 and the two CAPRI datasets were AP-PISA and CP-PIE.

DL methods performed significantly better than classical methods for the remaining four datasets, namely, BM4, Dockground, PDB-2023, and MaSIF-test. PIsToN had the highest AUC value for three of the four datasets, while GNN-DOVE outperformed its competitors for Dockground. We note that GNN-DOVE applied 4-fold cross-validation on this dataset for its training, validation, and test parts. Scores obtained from each of the four test sets were used for a fair comparison. For PDB-2023 and MaSIF-test dataset, PIsToN had significantly better performance than its closest competitor, dMaSIF, with AUC values of 93.86 and 91.83 for PDB-2023 and 93.55 and 89.90 for MaSIF-test dataset, respectively. Barring the CAPRI datasets, we conclude that DL-based methods outperformed classical methods on five of the seven datasets.

### Success Rate Measures

The second metric for evaluating tools is *success rate*, which measures how often a docking method provides at least one model of acceptable quality within its top 1, top 10, top 100, or top N predicted poses. A higher success rate can impact the success of drug discovery efforts.

The success rates of all eleven scoring functions on all datasets are shown in Table 1. For almost all datasets, DL-based methods outperformed the competing classical methods. For CAPRI score refined dataset, BM4, and PDB-2023 datasets, PIsToN had the most number of top ranking performances for top 1, top 10, top 25, top 100, and top 200 predictions. In the CAPRI score dataset, DL-based methods showed better or equal performance compared to others, where DeepRank-GNN and dMaSIF were the best for top 1 (23%), dMaSIF, PIsToN, CP-PIE, and SIPPER were altogether the best for top 10 (46%), and for top 100 and top 200 dMaSIF and PIsToN outperformed others with 76% and 92% success, respectively. For the BM5 dataset, both DL and classical approaches had comparable performances. DeepRank-GNN and CP-PIE were the best for all topN predictions, even reaching 100% success in many cases. For the Dockground dataset, GNN-DOVE had the highest performance for all topN predictions. For the top 100 and 200, all other DL methods and some classical methods (CP-PIE, ZRANK2, and SIPPER) had 100% success in ranking poses.

**Table 1.** Success rates of all eleven scoring functions on six different datasets. The top four methods are DL-based, and the rest are classical methods (details in Sec. 2)

| Dataset | Method | top1 | top10 | top25 | top100 | top200 |
| --- | --- | --- | --- | --- | --- | --- |
| **CAPRI**<br>**Score** | PIsToN | 15 | **46** | 69 | 69 | **92** |
|  | dMaSIF | **23** | **46** | 61 | **76** | 84 |
|  | DeepRank-GNN | **23** | 38 | 53 | 69 | 84 |
|  | GNN-DOVE | 0 | 15 | 30 | 69 | 69 |
|  | FireDock | 0 | 0 | 7 | 30 | 46 |
|  | AP-PISA | 0 | 0 | **70** | 53 | 53 |
|  | CP-PIE | 15 | **46** | 53 | 69 | 76 |
|  | PyDock | 0 | 0 | 0 | 15 | 46 |
|  | ZRANK2 | 7 | 30 | 46 | 69 | 69 |
|  | RosettaDock | 7 | 30 | 46 | 69 | 69 |
|  | SIPPER | 7 | **46** | 53 | 69 | 84 |
| **CAPRI**<br>**Score**<br>**Refined** | PIsToN | **38** | **69** | **76** | **76** | **100** |
|  | dMaSIF | 15 | 38 | 53 | **76** | 92 |
|  | DeepRank-GNN | 23 | 46 | 53 | 69 | 84 |
|  | GNN-DOVE | 15 | 61 | 69 | **76** | **100** |
|  | FireDock | 0 | 0 | 7 | 23 | 46 |
|  | AP-PISA | 0 | 0 | 7 | 46 | 53 |
|  | CP-PIE | 23 | 46 | 69 | 69 | 84 |
|  | PyDock | 0 | 0 | 0 | 7 | 30 |
|  | ZRANK2 | 7 | 38 | 53 | 61 | 61 |
|  | RosettaDock | 7 | 38 | 53 | 69 | 76 |
|  | SIPPER | 7 | 46 | 53 | 69 | 76 |
| **BM4** | PIsToN | **68** | **89** | **89** | 94 | **100** |
|  | dMaSIF | 47 | 68 | **89** | 94 | **100** |
|  | DeepRank-GNN | 21 | 73 | 73 | 89 | 94 |
|  | GNN-DOVE | 21 | 68 | 84 | **100** | **100** |
|  | FireDock | 26 | 84 | **89** | 89 | 94 |
|  | AP-PISA | 26 | 78 | **89** | 94 | 94 |
|  | CP-PIE | 52 | 78 | 84 | **100** | **100** |
|  | PyDock | 15 | 63 | 84 | 94 | 94 |
|  | ZRANK2 | 42 | 78 | **89** | 94 | 94 |
|  | RosettaDock | 42 | 73 | **89** | 94 | 94 |
|  | SIPPER | 47 | 63 | 68 | 78 | 94 |
| **BM5** | PIsToN | 66 | 93 | **100** | **100** | **100** |
|  | dMaSIF | 40 | 80 | **100** | **100** | **100** |
|  | DeepRank-GNN | **93** | **100** | **100** | **100** | **100** |
|  | GNN-DOVE | 6 | 26 | 33 | 53 | 73 |
|  | FireDock | 0 | 0 | 0 | 6 | 13 |
|  | AP-PISA | 0 | 6 | 6 | 13 | 26 |
|  | CP-PIE | **93** | **100** | **100** | **100** | **100** |
|  | PyDock | 0 | 0 | 0 | 6 | 6 |
|  | ZRANK2 | 0 | 0 | 6 | 6 | 20 |
|  | RosettaDock | 13 | 46 | 53 | 66 | 80 |
|  | SIPPER | 60 | 93 | 93 | **100** | **100** |
| **Dockground** | PIsToN | 15 | 55 | 81 | **100** | **100** |
|  | dMaSIF | 34 | 75 | 87 | **100** | **100** |
|  | DeepRank-GNN | 37 | 53 | 72 | **100** | **100** |
|  | GNN-DOVE | **70** | **91** | **94** | **100** | **100** |
|  | FireDock | 1 | 18 | 48 | 98 | **100** |
|  | AP-PISA | 1 | 6 | 17 | 98 | **100** |
|  | CP-PIE | 22 | 56 | 75 | **100** | **100** |
|  | PyDock | 0 | 8 | 20 | 93 | **100** |
|  | ZRANK2 | 5 | 25 | 53 | **100** | **100** |
|  | RosettaDock | 3 | 31 | 56 | 96 | **100** |
|  | SIPPER | 43 | 72 | 89 | **100** | **100** |
| **PDB**<br>**2023** | PIsToN | **88** | **96** | **98** | **100** | **100** |
|  | dMaSIF | 51 | 84 | 94 | **100** | **100** |
|  | DeepRank-GNN | 36 | 48 | 67 | **100** | **100** |
|  | GNN-DOVE | 3 | 19 | 34 | **100** | **100** |
|  | FireDock | 0 | 3 | 11 | **100** | **100** |
|  | AP-PISA | 0 | 1 | 5 | **100** | **100** |
|  | CP-PIE | 76 | 92 | 96 | **100** | **100** |
|  | PyDock | 0 | 0 | 3 | **100** | **100** |
|  | ZRANK2 | 0 | 11 | 23 | **100** | **100** |
|  | RosettaDock | 3 | 17 | 32 | **100** | **100** |
|  | SIPPER | 50 | 75 | 84 | **100** | **100** |

Bold values indicate the best value for each column.

### Comparing Run Times

The running time of each scoring function was also evaluated, and the results are shown in Figure 3. For this set of experiments, instead of running all the datasets, we subsampled the CAPRI score dataset, which is the most diverse of the datasets from Section 3, to randomly select one correct and one acceptable docking model for each complex, resulting in a smaller dataset with 26 docking models for 13 complexes. The average run times from these experiments are thus averaged over complexes from different tools. As shown in Figure 3, dMaSIF and GNN-DOVE, both DL-based methods, exhibit impressive efficiency, with average runtimes of 3 seconds and 7 seconds, respectively, surpassing the speed of all the classical methods. dMaSIF represents surface atoms as point clouds, calculating features on the fly without pre-computation, and this makes it faster than its competitors. We note that PIsToN and DeepRank-GNN have a preprocessing component included in the running time calculation. If preprocessing time is discounted, PIsToN requires 0.04 seconds, while DeepRank-GNN requires 6.3 seconds. Furthermore, since all classical tools were run on the CCharPPI server, their running times are approximate. The results show that DL-based methods are computationally more efficient than the classical methods.

**Fig. 3.**
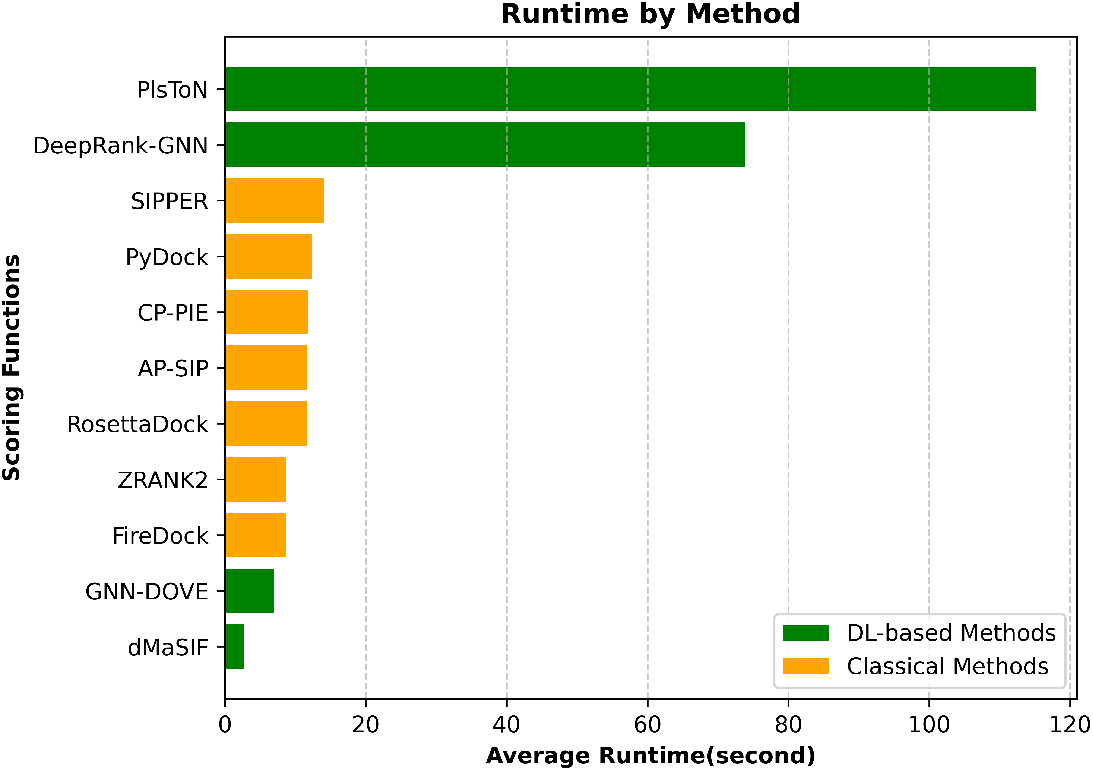
Average runtime of all eleven scoring functions on a dataset subsampled from the CAPRI Score dataset

## 5 Discussion and Conclusions

In this study, we conducted a comprehensive comparison between state-of-the-art DL-based and classical scoring functions. We used various popular datasets of different sizes. Among all classical methods, AP-PISA emerged as the top-performing method across all eleven datasets. Unlike other classical methods that mainly consider potentials from the residue level, AP-PISA re-ranks refined docking models using a combination of atomic and residue potentials, which helps to identify near-native poses correctly. On the other hand, AP-PISA relies solely on potential features, but incorporating energy terms can enhance performance by leveraging empirical and knowledge-based features. Overall, according to the AUC metric, CP-PIE was the second-best classical method and the best in ranking power. CP-PIE efficiently filters unlikely docking poses before performing computationally intensive scoring calculations. It discards misdocked structures by measuring the overlap area between the proteins in each docking model. One conclusion that can be drawn is that the refinement stage, which involves deleting far-away poses from the correct poses, is crucial for accurately scoring docking models.

In general, PIsToN and dMaSIF performed better than their DL-based competitors regarding both AUC and success rate metrics across all datasets. PIsToN utilized the strength of Vision Transformers [98] and contrastive training technique [110] to incorporate all chemical, spatial, and energy features, resulting in superior performance compared to other methods. However, PIsToN lagged behind its rival dMaSIF in terms of running time measurement due to its preprocessing step. PIsToN precomputed meshes for the surface of the two proteins involved in generating the complex to capture topological relationships and the geometry of their surfaces, enabling it to be more accurate, but this process is computationally intensive. It may be beneficial for PIsToN to improve the preprocessing time by changing the surface representation to use non-Euclidean structural data like point clouds or graphs instead of meshes. On the other hand, dMaSIF has a very fast running time. It represents the surface atoms as a set of point clouds, providing a flexible and simpler representation that can easily handle complex structures. For each atom, it only considers the atom type and its coordinates to calculate chemical features, avoiding the computation of physico-chemical features and significantly improving the runtime. However, dMaSIF estimates the chemical features based solely on the atom types and coordinates, failing to take into account the impact of other types of features in binding, such as energy terms, which can compromise accuracy.

Our study has demonstrated that DL-based techniques have proven to be significantly superior to classical methods for five out of seven datasets when measured by the AUC metric. Classical methods have limitations due to their inability to account for the 3D features and nonlinear relations between energy terms. This drawback affects their performance. Moreover, deep learning-based methods have consistently shown superior performance across all eleven datasets when evaluated using the success rate metric. This is particularly important in fields such as drug discovery and protein engineering, where accurate identification of protein configurations can lead to fewer experiments and a decrease in experimental costs, thereby providing more reliability and cost efficiency in such applications.

In this study, we only evaluated rigid-body methods since most researchers have proposed scoring functions based on these methods. However, it might be worth exploring potential future directions to introduce scoring functions that also consider the flexibility of proteins during complex formation. Another potential path for future work could be to develop approaches for scoring multi-chain complexes and generating appropriate datasets to evaluate their performance.

## Data Abvailability

The datasets used in this study can be found at the following links: CAPRI Score: http://cb.iri.univ-lille1.fr/Users/lensink/Score_set/; CAPRI Score Refined and BM5: https://data.sbgrid.org/dataset/843/; 15 chosen complexes from BM5: https://doi.org/10.5281/zenodo.10442444; Dockground: http://dockground.compbio.ku.edu/downloads/unbound/decoy/decoys1.0.zip; BM4: http://zlab.umassmed.edu/benchmark/; 19 chosen complexes from BM4 https://doi.org/10.5281/zenodo.10442444; PDB-2023: https://zenodo.org/record/7948337/files/PDB_2023.tar.gz; and MaSIF-test: https://zenodo.org/record/7948337/files/masif_test.tar.gz.

## Acknowledgements

We thank the members of the Bioinformatics Research Group for valuable feedback. This work was partially funded by NSF grants CNS-203734 and OAC-2118329, a GAANN award from the Department of Education, and the Knight Foundation School of Computing and Information Sciences.

