## Supplementary Material for "A Comprehensive Survey of Scoring Functions for Protein Docking Models"

**Table S1.** Classification performance of scoring functions on seven datasets. The Columns correspond to the Area Under the Receiver Operating Characteristic curve (AUC ROC), Average Precision (AP), Balanced Accuracy (BA), and F1 scores, respectively. Bold values indicate the best value for each column.

| Dataset | Method | AUC ROC | AP | BA | F1 | Precision | Recall |
|---|---|---|---|---|---|---|---|
| **CAPRI Score** | PIsToN | 77.48 | 24.36 | **75.58** | 40.35 | 26.47 | 84.92 |
| | dMaSIF | 73.60 | 12.48 | 49.84 | 22.06 | 12.48 | 95.04 |
| | DeepRank-GNN | 66.42 | 12.41 | 40.46 | 7.61 | 6.51 | 19.56 |
| | GNN-DOVE | 56.70 | 13.10 | 52.56 | 23.12 | 13.12 | **97.16** |
| | FireDock | 67.20 | 17.23 | 64.88 | 29.81 | 18.11 | 84.29 |
| | AP-PISA | **77.64** | **24.49** | 73.74 | **41.40** | 28.83 | 73.40 |
| | CP-PIE | 74.34 | 12.77 | 51.14 | 22.50 | 12.79 | 93.59 |
| | PyDock | 70.36 | 20.68 | 68.12 | 36.34 | 25.75 | 61.69 |
| | ZRANK2 | 66.00 | 21.62 | 67.15 | 38.07 | **30.36** | 51.04 |
| | RosettaDock | 61.98 | 18.44 | 64.88 | 32.66 | 22.76 | 57.83 |
| | SIPPER | 55.43 | 14.43 | 56.88 | 24.82 | 15.94 | 56.00 |
| **CAPRI Score Refined** | PIsToN | 81.25 | 24.86 | 73.24 | 42.21 | 30.37 | 69.16 |
| | dMaSIF | 78.53 | 12.33 | 49.14 | 21.89 | 12.33 | 97.69 |
| | DeepRank-GNN | 76.01 | 12.53 | 50.07 | 22.23 | 12.53 | **99.81** |
| | GNN-DOVE | 70.99 | 10.35 | 34.90 | 14.06 | 8.18 | 49.88 |
| | FireDock | 72.79 | 19.00 | 67.50 | 33.01 | 21.17 | 74.89 |
| | AP-PISA | **84.87** | **30.18** | **75.36** | **49.29** | **39.80** | 67.72 |
| | CP-PIE | 81.29 | 12.31 | 49.05 | 21.84 | 12.31 | 97.16 |
| | PyDock | 75.28 | 25.03 | 70.49 | 42.98 | 34.93 | 55.86 |
| | ZRANK2 | 74.09 | 25.68 | 69.57 | 43.76 | 38.46 | 50.75 |
| | RosettaDock | 63.95 | 18.94 | 64.75 | 33.64 | 24.88 | 51.90 |
| | SIPPER | 53.03 | 13.89 | 55.25 | 23.62 | 14.97 | 55.95 |
| **BM4** | PIsToN | **67.45** | **36.46** | **62.71** | **43.67** | **53.75** | 36.77 |
| | dMaSIF | 62.50 | 26.27 | 49.64 | 41.44 | 26.27 | **98.16** |
| | DeepRank-GNN | 59.59 | 26.41 | 50.00 | 41.58 | 26.41 | 97.76 |
| | GNN-DOVE | 61.58 | 25.11 | 46.47 | 38.04 | 24.82 | 81.37 |
| | FireDock | 57.82 | 28.59 | 54.28 | 30.18 | 34.53 | 26.81 |
| | AP-PISA | 59.21 | 25.36 | 47.19 | 39.40 | 25.25 | 89.74 |
| | CP-PIE | 59.19 | 24.23 | 43.37 | 31.40 | 22.30 | 53.01 |
| | PyDock | 58.31 | 30.16 | 57.71 | 41.65 | 33.10 | 56.15 |
| | ZRANK2 | 55.85 | 24.97 | 45.97 | 35.24 | 24.20 | 64.82 |
| | RosettaDock | 57.37 | 24.71 | 44.64 | 26.55 | 21.51 | 34.68 |
| | SIPPER | 59.87 | 24.50 | 42.28 | 18.21 | 16.80 | 19.88 |

Bold values indicate the best value for each column.

| Dataset | Method | AUC ROC | AP | BA | F1 | Precision | Recall |
|---|---|---|---|---|---|---|---|
| **BM5** | PIsToN | 91.84 | 23.64 | 74.90 | 45.14 | 38.16 | 55.24 |
| | dMaSIF | 87.89 | 4.39 | 27.24 | 5.86 | 3.11 | 51.05 |
| | DeepRank-GNN | 95.97 | 5.72 | 49.66 | 0.0 | 0.0 | 0.0 |
| | GNN-DOVE | 60.76 | 9.00 | 53.69 | 13.57 | 47.22 | 7.93 |
| | FireDock | 95.41 | **43.12** | **85.00** | **63.96** | 56.65 | **74.43** |
| | AP-PISA | 91.62 | 33.45 | 72.24 | 54.20 | 66.11 | 45.92 |
| | CP-PIE | **97.16** | 5.72 | 49.37 | 0.0 | 0.0 | 0.0 |
| | PyDock | 88.98 | 19.61 | 67.05 | 39.90 | 43.21 | 37.06 |
| | ZRANK2 | 90.37 | 29.20 | 71.00 | 50.40 | 59.31 | 43.82 |
| | RosettaDock | 72.26 | 40.56 | 77.17 | 61.38 | **67.99** | 55.94 |
| | SIPPER | 87.70 | 4.86 | 18.09 | 2.67 | 1.43 | 20.05 |
| **Dockground** | PIsToN | 68.42 | 10.71 | 60.40 | 22.13 | 15.62 | 37.92 |
| | dMaSIF | 79.94 | 6.33 | 26.10 | 5.71 | 3.15 | 31.46 |
| | DeepRank-GNN | 59.76 | 7.75 | 50.21 | 3.93 | 9.16 | 25.00 |
| | GNN-DOVE | **83.87** | 6.24 | 23.61 | 5.53 | 3.03 | 31.46 |
| | FireDock | 56.08 | 8.50 | 54.82 | 15.69 | 8.82 | 70.83 |
| | AP-PISA | 79.19 | **18.14** | **64.77** | **36.08** | **38.23** | 34.17 |
| | CP-PIE | 72.26 | 6.50 | 38.67 | 10.65 | 5.81 | 63.75 |
| | PyDock | 77.17 | 13.71 | 64.10 | 28.99 | 23.01 | 39.17 |
| | ZRANK2 | 64.18 | 10.78 | 58.76 | 22.35 | 18.93 | 27.29 |
| | RosettaDock | 61.62 | 9.41 | 57.17 | 18.50 | 12.80 | 33.33 |
| | SIPPER | 79.60 | 7.75 | 50.28 | 14.39 | 7.75 | **99.17** |
| **PDB 2023** | PIsToN | **93.86** | **45.85** | **78.21** | **66.05** | **78.89** | 56.80 |
| | dMaSIF | 91.83 | 1.85 | 23.46 | 2.11 | 1.08 | 42.40 |
| | DeepRank-GNN | 62.65 | 2.43 | 50.36 | 3.69 | 2.76 | 56.00 |
| | GNN-DOVE | 53.23 | 2.55 | 52.59 | 5.16 | 2.76 | 40.00 |
| | FireDock | 87.98 | 16.68 | 67.39 | 38.79 | 42.06 | 36.00 |
| | AP-PISA | 89.19 | 21.68 | 68.02 | 44.02 | 54.76 | 36.80 |
| | CP-PIE | 88.81 | 2.39 | 46.71 | 0.40 | 0.27 | 8.00 |
| | PyDock | 87.64 | 10.48 | 59.30 | 26.82 | 44.44 | 19.20 |
| | ZRANK2 | 73.22 | 23.31 | 66.19 | 43.85 | 66.13 | 32.80 |
| | RosettaDock | 65.04 | 20.67 | 66.48 | 42.21 | 56.76 | 33.60 |
| | SIPPER | 76.25 | 2.39 | 49.52 | 4.65 | 2.39 | **93.60** |
| **MaSIF test** | PIsToN | **93.55** | **81.08** | **85.43** | **85.97** | **87.72** | 82.40 |
| | dMaSIF | 89.90 | 45.55 | 19.60 | 13.52 | 14.63 | 12.57 |
| | DeepRank-GNN | 69.24 | 47.92 | 45.64 | 62.05 | 47.66 | 88.90 |
| | GNN-DOVE | 55.71 | 50.85 | 51.63 | 38.76 | 52.81 | 30.62 |
| | FireDock | 77.82 | 48.33 | 46.52 | 62.79 | 48.15 | 90.24 |
| | AP-PISA | 76.43 | 49.13 | 48.22 | 64.32 | 49.07 | **93.34** |
| | CP-PIE | 76.98 | 63.55 | 70.56 | 74.29 | 65.94 | 85.06 |
| | PyDock | 74.73 | 48.53 | 46.97 | 63.40 | 48.40 | 91.86 |
| | ZRANK2 | 74.66 | 49.06 | 48.08 | 64.11 | 48.98 | 92.75 |
| | RosettaDock | 71.17 | 49.13 | 48.22 | 63.99 | 49.05 | 92.01 |
| | SIPPER | 66.17 | 56.01 | 60.43 | 66.63 | 57.61 | 78.99 |

Bold values indicate the best value for each column.

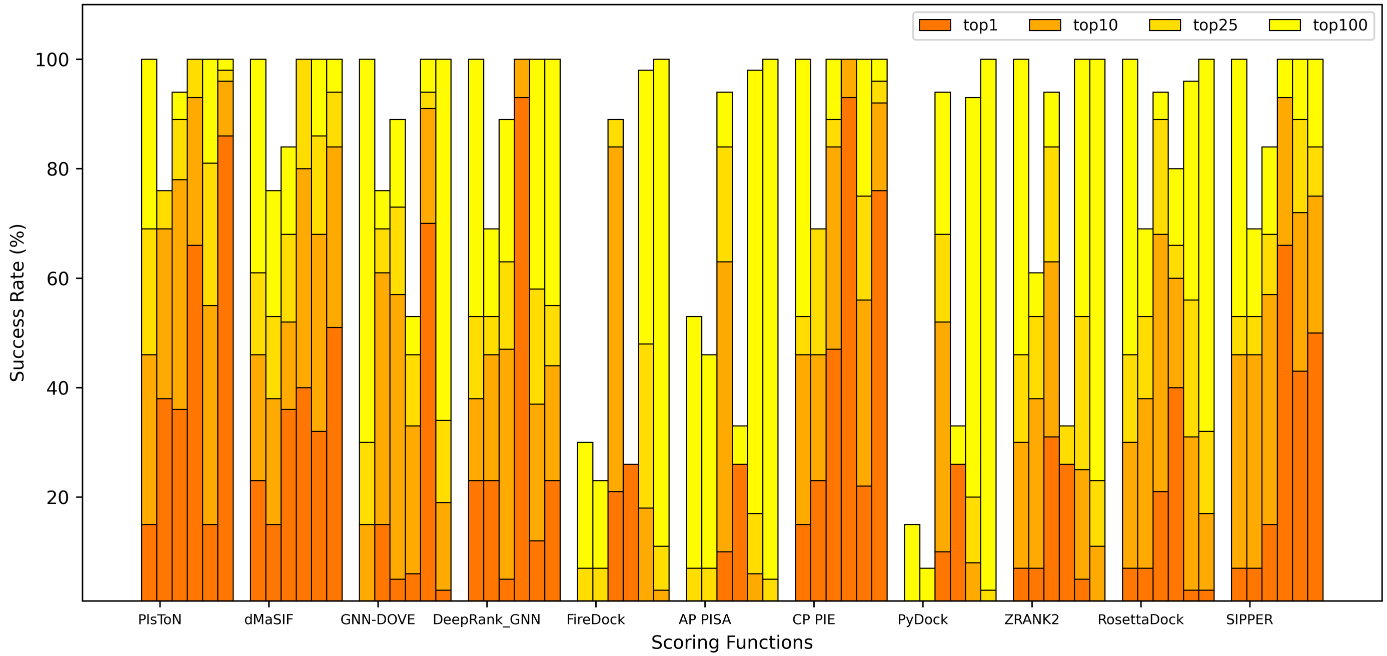


**Fig. S1.** The accumulative percentage of success rates of scoring functions. For each method, the columns correspond to datasets (from left to right: CAPRI Score, CAPRI Score Refined, BM4, BM5, Dockground, and PDB-2023). The colors correspond to the top 1, top 10, top 25, and top 100.
